# GelMetrics: An Algorithm for Analyzing the Dynamics of Gel-Like Phase-Separated Condensates

**DOI:** 10.1101/2024.07.03.601857

**Authors:** Amit Nelkin, Naor Granik, Roee Amit

## Abstract

Cellular compartments and organelles are essential for the spatial organization of biological matter. Recently, membraneless organelles like paraspeckles, stress granules, and Cajal bodies have garnered significant scientific interest due to their lack of membrane boundaries and crucial cellular functions. These organelles self-assemble through phase separation, a process in which a homogeneous solution separates into distinct phases. The phases most commonly encountered in cells are liquids and gels. Although various microscopy techniques exist to study phase-separated compartments, they are often inadequate for investigating the dynamics of gel-like condensates, where molecular motion occurs over tens of minutes rather than seconds. Here, we introduce the GelMetrics algorithm to quantitatively measure the dynamics of gel-like phase-separated structures by tracking their fluorescence signals over extended durations. First, we show using simulations that this method can identify molecular motion amidst measurement noise and estimate biophysical parameters such as the number of molecules and the average fluorescence of a single molecule exiting a condensate, providing insights into the dynamics of phase-separated organelles. Second, we validate our approach on data from synthetic RNA-protein (sRNP) granules in *E. coli* and *in vitro*, establishing its applicability both *in vitro* and *in vivo*. Finally, we demonstrate the biological meaning of our algorithm’s output by quantitatively determining how the presence of an intrinsically disordered region causes sRNP granules to exhibit a more brittle phase-behavior. Thus, not only does the proposed method fill a gap in analysis methods for gel-like condensates, but it also elucidates biophysical mechanisms, improving our understanding of both synthetic and natural systems.

## I. INTRODUCTION

Cellular compartments and organelles play a crucial role in the structural organization of biological matter. Well-established compartments such as the nucleus or Golgi apparatus are distinguished by the presence of a lipid membrane boundary that separates them from the surrounding cellular environment. In addition to these conventional membrane-bound organelles, there exists another category of cellular compartments known as membraneless organelles. This category includes compartments such as paraspeckles, stress granules, Cajal bodies, etc [1,2]. These cellular entities are unique in that they remain distinct from the cellular environment despite the absence of a membrane, while performing various functions. Recent research suggests that these membraneless organelles, often characterized as supramolecular assemblies of proteins and RNA molecules, self-assemble through the process of phase separation [3–6].

Phase separation is the process by which a homogeneous solution separates into multiple distinct phases, each with unique compositions and properties. Typically, phase separation processes are classified by the different material states of the resulting phases (e.g., liquid, gel, glass), which can lead to various phase transitions (for example, a liquid-solid transition indicates a transition from a liquid environment into a solid-like compartment). The transitions commonly encountered in cellular biology are liquid-liquid and liquid-gel. However, determining the characteristics and driving forces in a phase separating system within a cellular environment remains a challenging task.

A primary method of investigation of naturally occurring membraneless organelles is *in vitro* reconstitution. This method entails incubation of individual components of condensates in specific conditions to test whether they phase separate *in vitro* [7–9]. Alternative methods attempt to probe the dynamics of membraneless organelles using microscopy. These include fluorescence recovery after photobleaching (FRAP), which is utilized to assess the liquidity of phase separated condensates, and fluorescence resonance energy transfer (FRET), which can measure interaction kinetics within condensates, including binding and unbinding kinetics. More recently, fluorescence correlation spectroscopy (FCS) has been utilized to investigate diffusivity at the level of individual molecules, assuming sufficient sparsity [10–12].

Although FRAP is perhaps the most extensively utilized fluorescence-based method, its applicability is limited to scenarios in which the condensate is predominantly liquid-like, in which case diffusion can be observed and measured. Alternatively, FRAP is useful when attempting to determine the material state itself, as fluorescence recovery, or lack thereof, can differentiate between liquid-like and solid-like condensates. Notably, FRAP is not always suitable when investigating dynamics in condensates with solid-like characteristics, specifically, gel-like and glass-like, where molecular motion exists but at a significantly slower rate [13–15].

Given the slower molecular dynamics of gel-like, glass-like, or other viscoelastic condensates, we propose an alternative approach for their study that does not rely on specialized microscopy methods. By tracking condensate fluorescence intensities over extended periods, we can address the issue of low diffusivity within the granules by analyzing the fluorescence signals themselves to extract instances of molecular motion at the phase boundary. The challenge of finding a weak or rare signal in a noisy environment is known as rare signal recovery. Basic techniques in this field, such as matched filtering and Bayesian analysis, utilize prior knowledge of the underlying signal to separate it from the noise. More advanced approaches, like subspace methods and empirical mode decomposition, provide robust tools for identifying weak signals. These methods and others were designed to tackle complex signals commonly encountered in communications and remote sensing (e.g., radar). As such, they might be less effective in the biological context described here.

In this work, we present a method to quantitatively measure the dynamics of phase-separated, gel-like granules called GelMetrics, by tracking their fluorescence signals over extended periods of time. We demonstrate on simulated signals that our GelMetrics algorithm can distinguish between different types of possible signals (constant, gradually changing, and bursty). Moreover, our algorithm successfully extracts important biophysical features of the granules from the signals, even in the presence of measurement noise. We then apply our algorithm to synthetic RNA-protein granules, where the protein component has an RNA binding moiety and either does or does not contain an intrinsically disordered region (IDR), and the RNA component has a variable valency. Our data analysis shows that granules containing proteins without an IDR exhibit gel-like diffusion behavior characterized by a steady-state rate of either release or absorption of single RNA molecules on the timescale of minutes, independent of valency. However, granules with a protein component containing an IDR element exhibit a more brittle dynamic behavior for low RNA valency, that is characterized by chunks of dense solid-like or glass-like granules (i.e. multiple protein-bound RNA molecules) that occasionally split o. ff from the main granule body. Consequently, we demonstrate that our signal analysis algorithm can be used as an additional tool for characterizing the biophysical properties of phase-separated, non-liquid condensates within cells.

## II. NUMERICAL MODEL

### A. Assumptions

We refer here to a scenario of a gel-like phase-separated, localized granule, formed by interactions among either fluorescent or fluorescently labeled components, residing within a dilute solution that can itself contain a low concentration of fluorescent molecules [Fig. 1]. Typically, at least one component is multivalent, enabling both inter- and intramolecular interactions that promotes granule formation. We assume an experimental setting wherein a field of view containing both highly fluorescent granules and a dark background is monitored for a total duration (T), with constant intervals (t) between fluorescence intensity measurements.

**FIG. 1.**
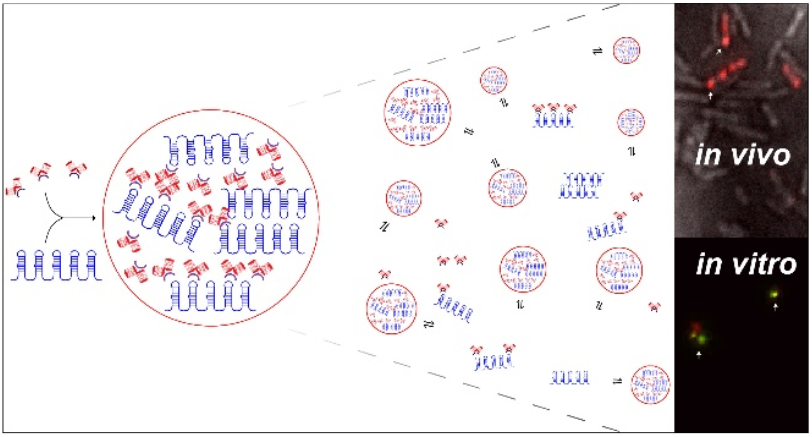
Formation of gel-like granules. Left: in our system, a synthetic long non-coding RNA (slncRNA) molecule encodes multiple repeats of a hairpin structure. These hairpins are specifically bound by an RNA-binding protein (RBP) fused to a fluorescent protein. Middle: these interactions drive phase separation, sequestering RNA and RBP molecules from the dilute solution while maintaining a slow rate of molecule exchange with the surrounding environment. Right: granules form both *in vivo* (appearing as fluorescent puncta at the cell poles) and *in vitro* (appearing as localized structures composed of fluorescent protein and fluorescently labeled synthetic RNA molecules).

We assume that the total fluorescence of the granule consists of three signal components: granule fluorescence, background fluorescence, and noise. Granule fluorescence represents the sum of fluorescence intensities of the molecules within the granule core, with occasional addition or subtraction of intensity resulting from fluorescent molecules entering or exiting the granule. Background fluorescence denotes the fluorescence intensity of the surrounding dilute phase, which contains similar components to the granule, albeit at a much lower concentration. Noise here refers to various sources that commonly affect fluorescence microscopy, including shot noise, electronic noise, and dark current noise [16]. The sum of these processes is assumed to be a symmetric, memory-less process (i.e., white noise). We further assume that background fluorescence changes slowly over time, in contrast to granule fluorescence, which depends on the dynamic insertion and shedding events occurring in the granule.

Based on these assumptions, we define a general model for the fluorescence intensity signal that includes both additive and exponential components, representing fluorescent molecules, and photobleaching, respectively. The model can be described as follows:

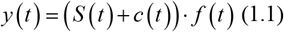

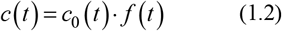

Where *y*(*t*) is the fluorescence signal measured experimentally, *S*(*t*) denotes the underlying granule signal, *c*(*t*) is the observed background signal, *c*_0_(*t*) is the underlying background signal, and *f*(*t*) is the photobleaching component, which affects both granule and background signals equally.

### B. Signal extraction

To find the underlying granule signal *S*(*t*), we assume that the background signal is a constant:

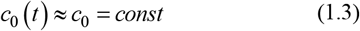

This leads to:

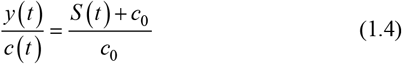

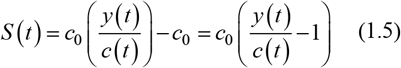

Notably, this model lacks the contribution of high-frequency noise effects stemming from physical measurements in a fluorescence microscope. These affect the measured granule signal *y*(*t*), and the measured background signal *c*(*t*). To reduce these effects, and since we do not require the background signal at high temporal resolution, we fit the measured background signal *c*(*t*) to a 3rd degree polynomial. This is done to capture the general trend of the signal (i.e., the photobleaching component) while eliminating fluctuations due to random noise. In contrast, the measured granule signal *y*(*t*), which contains biologically relevant data, is filtered with a moving average filter with a window span dependent on experimental considerations.

### C. Identifying signal bursts

A hallmark of phase-separated compartments is the movement of macromolecules across the phase boundary in either direction [17,18]. In terms of fluorescence intensity, such movement should manifest as a change in signal intensity, lasting over multiple time points and resulting in a higher or lower stable signal level. We term such shifts in the signal as “signal bursts”. To identify such shifts in the baseline fluorescence intensity, we apply a moving-average filter to smooth the data as described in the signal extraction section. This operation transforms the fluctuations of the smoothed noisy signal near the bursts towards a gradual increase or decrease in the signal [Fig. 2(a)]. Random fluctuations that do not establish a new baseline level are unlikely to produce a gradual and continuous increase or decrease over multiple time points. We note that this operation alone can mask biologically relevant events that occur on very short timescales, rendering this algorithm less effective for liquid-like condensates with fast molecular motion.

**FIG. 2.**
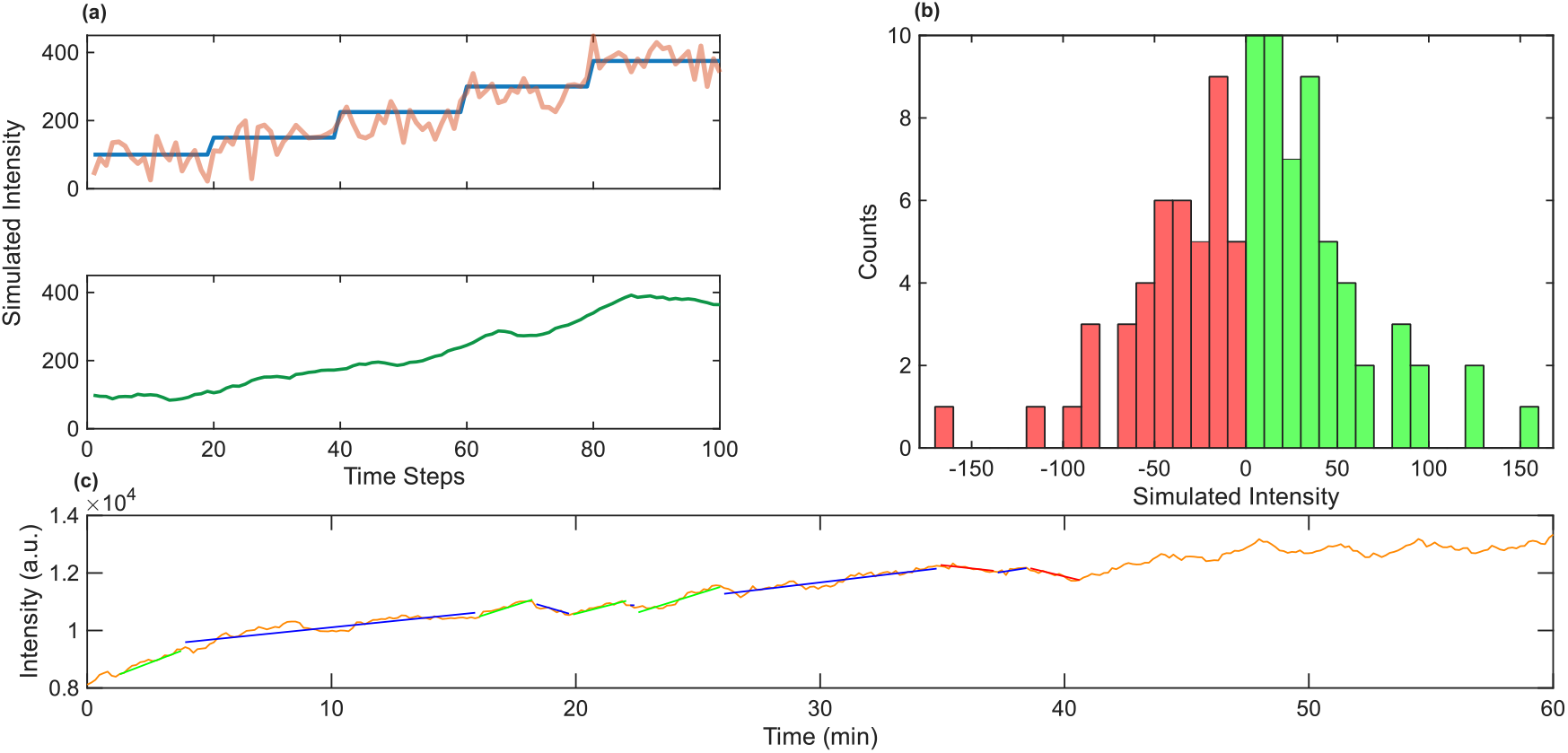
Identification of burst events. (a) Top: simulated step signal (blue) with added white Gaussian noise (orange). Bottom: noisy signal after application of a moving-average filter, demonstrating continuous changes in the signal lasting for several time points. (b) Intensity difference distribution for the signal presented in panel (a). Such distribution can be used to compute the likelihood of observing an instantaneous signal increase event at a specific time-point. (c) Sample experimental signal (orange) overlaid with markers indicating identified segments: positive bursts (green), unclassified segments (blue), and negative bursts (red).

Following this filtering step, we search for contiguous segments of gradual increase or decrease, and record only those with a probability for occurrence of 1 in 1000 or less under the null hypothesis of white noise. To translate this probability into a computational threshold, we first compute the intensity difference distribution for each trace. This distribution is obtained by collecting all of the instantaneous differences in the signal Δ*S*(*t*_i_) *= S*(*t*_i_) *-S*(*t*_i-1_) and binning them [Fig. 2(b)]. For a given trace, the likelihood of observing an instantaneous signal increase event at a specific time point *t*_*i*_ can be computed as follows:

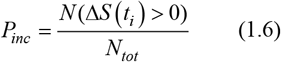

Where *N*(Δ*S*(*t*_*i*_) > 0) and *N*_*tot*_ denote the number of increasing instantaneous events, and the total number of events in the trace, respectively. Similarly, the likelihood of observing decreasing instantaneous events is defined as:

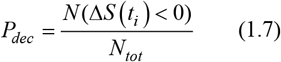

These likelihoods allow us to compute the number of consecutive instantaneous signal shift events *m* that satisfy our probability threshold for a significant signal burst event, as follows:

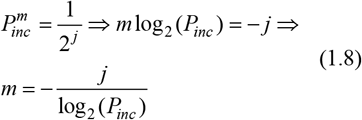

Here, *j* is a stringency parameter which allows us to control the threshold probability. For example, a value of *j*=10 would correspond to a probability of 1/1024, which is the maximal probability that we allow. This freedom of stringency is required since different systems will have different levels of noise, resulting in more false-positive events being detected. The value of *m* that satisfies the probability threshold is calculated per signal, and an analogous threshold *n* is calculated for decrements in the signal, with *n* typically in the range [*m*-1, *m*+1]. We refer to these *m* (or more) consecutive monotonous frames as a single burst. More generally, we define a burst as a set of time points equivalent to *L* consecutive monotonous frames where *L* > min(*m,n*). For each burst, we record the amplitude *I*_*burst*_, defined as the difference between the signal values at the final and first time points of the burst. Additionally, we record the time between successive events of the same type (e.g., the time between two positive burst events).

Using this definition of bursts, we can identify segments within the signal trace corresponding to signal bursts. Segments for which *I*_*burst*_ > 0 and *n* ≤ *L* ≤ *m*, or segments for which *I*_*burst*_ < 0 and *m* ≤ *L* ≤ *n* are considered unclassified. These are typically signal elements with a noise profile that does not allow us to make a classification. In Fig. 2(c), we mark the classifications on a sample trace with positive bursts, negative bursts, and non-classified events in green, red, and blue, respectively. Our segment analysis is confined to the range between the first and last significant segments identified in each signal, as we cannot classify sections that extend beyond the observed trace, as we have no recorded time difference to the following segment.

### D. Estimating signal parameters

Our model posits that a varying number of discrete molecules participate in a burst event (either positive or negative), hence a Poisson distribution is the most appropriate for describing the data. However, since we cannot directly infer the fluorescence intensity of a single macromolecule, we instead fit the burst amplitude distribution to a modified Poisson function of the form:

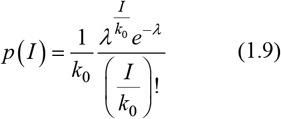

Here, *I* is the observed fluorescence amplitude of the burst, *λ* is the Poisson parameter that corresponds to the average number of molecules traversing the phase boundary per burst, and *k*_*0*_ is a fitting parameter whose value corresponds to the fluorescence intensity of a single molecule. For each pair of *λ* and *k*_*0*_ values, we calculate the theoretical Poisson distribution [Fig. 3] and evaluate its deviation from the experimental amplitude distribution. Over the course of our simulations, we observed that the algorithm performed insufficiently at first (Appendix A-C). Since the fit deviates from the sample data in low signal intensity values, which are of less significance, we decided to reduce the low-amplitude bias. First, we neglect low-intensity signals by left-truncating the amplitude data at the 15^th^ percentile threshold. Second, we calculate a weighted score (*WS*) to assess the deviation of the theoretical Poisson, combining three metrics: the mean squared error (MSE), the quantile-quantile (QQ) plot root mean squared error (RMSE), and the Kolmogorov–Smirnov statistic (KS):

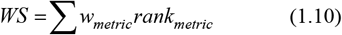

Where *w*_*metric*_ is the weight given to each metric and *rank*_*metric*_ is the rank-normalized metric score. We then present the estimates for *λ*=1,2,3,4,5 together with the corresponding truncated empirical QQ-plots and cumulative density function plots (CDF) [Fig. 12].

**FIG. 3.**
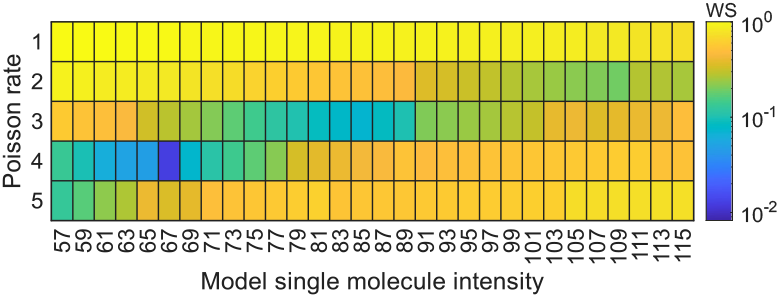
Poisson fitting example results. The heatmap depicts *WS* values as a function of *λ* and *k*_*0*_ values for simulated data with *λ*=4, later presented in Figs. 5(c) and 5(f).

## III. DIFFERENTIATING BETWEEN DIFFERENT SETS OF SIMULATED DATA

As an initial evaluation of our algorithm, we simulated three distinct types of base signals. For each simulation category, we generated and analyzed 1000 signals consisting of 360 time points to align with our experimental setup. The three signal types are: (1) signals featuring multiple burst events, emulating the characteristic behavior of phase-separated condensates, namely molecules transitioning between dense and dilute phases; (2) constant flat signals, representing a stationary observable akin to a phase-separated solid, (e.g., a glass phase with arrested dynamics); and (3) gradually increasing signals, simulating a continuous flux of fluorescent molecules moving unidirectionally. In all simulations, we incorporated two noise components according to our noise model: white Gaussian noise with a peak-to-peak amplitude of 40 (a.u.), matching the value estimated from experimental traces in our hands, and an exponential component, simulating photobleaching [Fig. 4].

**FIG. 4.**
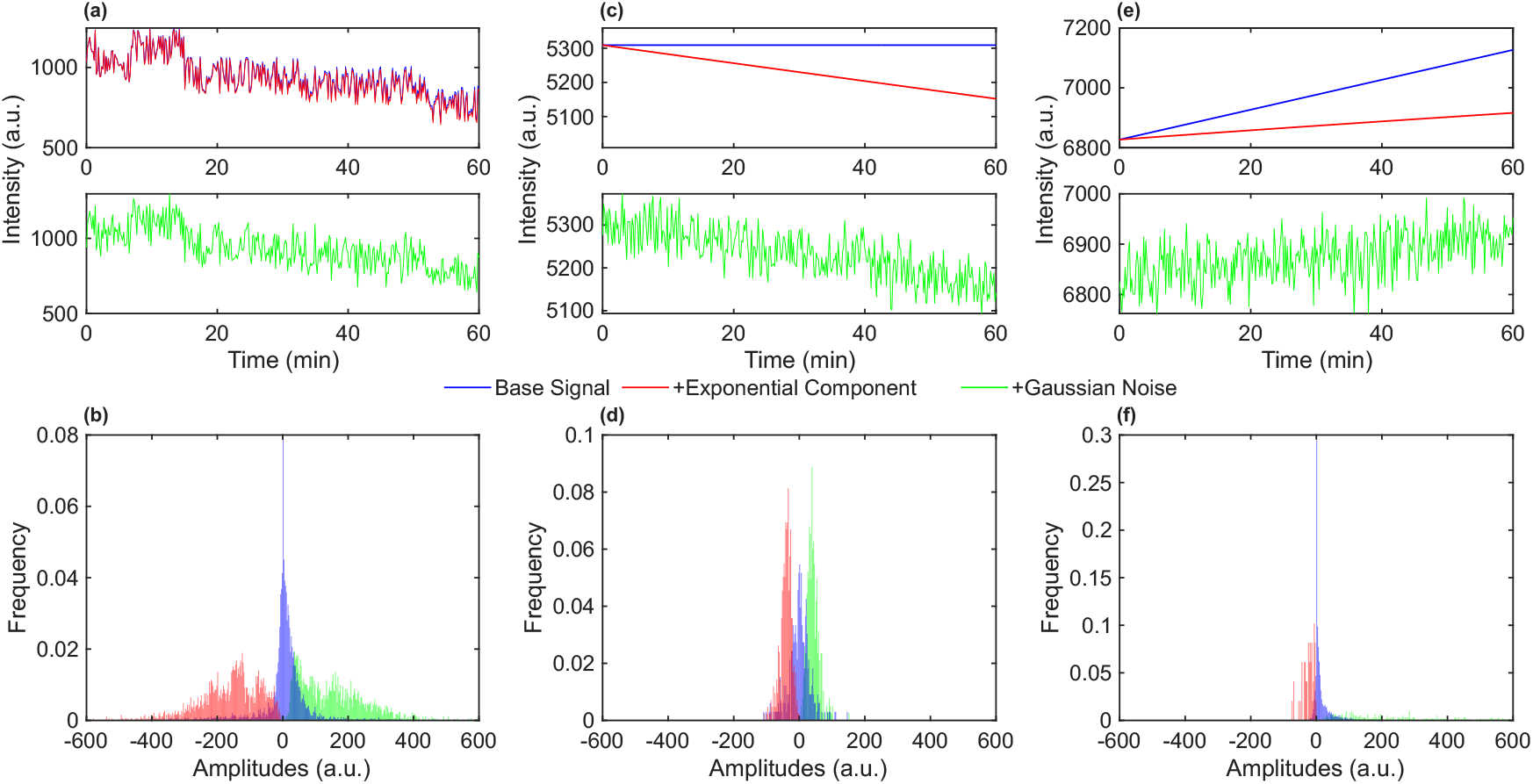
Differentiation between different types of signals. (a) Simulated signal with burst events (blue), with an added photobleaching component (red), and added Gaussian noise component (green). (b) Amplitude distributions of burst events identified from 1000 signals containing intermittent bursts. (c) Simulated constant signal (blue), with an added photobleaching component (red), and added Gaussian noise component (green). (d) Amplitude distributions of burst events identified from 1000 constant signals. (e) Simulated gradually ascending signal (blue), with an added photobleaching component (red), and added Gaussian noise component (green). (f) Amplitude distributions of burst events identified from 1000 sloped signals.

Prior to analysis, we first calibrated our algorithm such that the constant signals would output roughly one identified event per 1000 time points. For this, we applied our burst detection algorithm on the flat constant signals with three values of stringency parameter *j* [see Eq. (1.8)], 10, 12, and 15, corresponding to probabilities of 1/1024, 1/4096, and 1/32,768. In accordance with the given parameters, our algorithm identified 4.8, 3.2, and 1.7 segments per 1000 time points. Thus, a threshold parameter of *j*=15, corresponding to a probability of 1/32,768, was determined to be optimal for the simulation parameters stated above. We first applied our burst-detection algorithm to 1000 signals containing randomly distributed, increasing, or decreasing instantaneous bursts with an amplitude threshold of 60*A* (a.u), with *A* being a random number generated from a Poisson distribution with *λ*=2 [Fig. 4(a)]. The resulting burst amplitudes lie in the range 0-600 (a.u.) and appear symmetric in nature, with relatively similar numbers of positive and negative bursts detected [Fig. 4(b)]. Analysis of the constant simulated signals [Figs. 4(c) and 3(d)] reveals once again a symmetric positive and negative amplitude distribution; however, a close examination reveals that the burst amplitude width is smaller by a factor of ~5 compared to the results of the bursty signals. To compute whether the number of burst events identified via our algorithm is statistically significant, we utilized a chi-squared test. Our algorithm identified 383 and 418 positive and negative bursts in 1000 constant signals, respectively. For 1000 simulated signals containing bursts, 1700 and 1849 positive and negative bursts were identified, respectively, yielding a p-value < 1e-5, indicating a statistically significant difference between constant and bursty signals. In addition, for the constant signal case, the total number of events observed is ~400 for both positive and negative events, which indicates roughly one event per 1000 time points, in accordance with our choice of *j*. For the gradually increasing signals [Figs. 4(e) and 4(f)], a negligible number of negative burst-like events was detected by our algorithm, with a pronounced bias towards positive events (~50 negative events vs. ~600 positive events). The scarcity of events can be explained by the positive bias in the signal, which results in a steep increase in the statistical threshold for event identification *m*.

Similar simulations with a decreasing signal show a mirror image of burst amplitude distribution (data not shown). Taken together, these results demonstrate that our algorithm can differentiate between different types of signals, each corresponding to a distinct underlying experimental case.

## IV. ESTIMATING SIMULATED SIGNAL PARAMETERS

The simulated bursty signals were generated with burst amplitudes that are a product of two values: a constant intensity *k*_*0*_, and a random number generated from an underlying Poisson distribution with a specified *λ* parameter. To examine whether we can correctly identify the signal parameters, we generated three sets of simulated signals with different *λ*_*sim*_ values: 1,2,4. We then fitted the extracted negative amplitude data (in absolute values) to our modified Poisson function. Fig. 5 displays the negative amplitude distributions for all three cases, demonstrating a visible change in distributions with higher *λ*_*sim*_ values [Figs. 45(a)-5(c)] The estimation results corresponding to all *λ* values used in the simulation are presented [Figs. 5(d)-5(f)]. During the fitting process, we empirically observed improved algorithm performance when some percentage of the highest-amplitude signals were excluded from the input data. Specifically, the top 10% of amplitudes were removed. We attribute this effect to the low-probability nature of extreme signal values, which can disproportionately influence the fit and bias it toward the distribution tail, contrary to the algorithm’s intended purpose of describing the bulk of the distribution. In all three scenarios, the algorithm managed to estimate a *k*_*0*_ value relatively close to the ground truth [60 (a.u.)] and yielded correct *λ*_*est*_ values, except for *λ*_*sim*_=2. For *λ*_*sim*_=1, the algorithm yielded *λ*_*est*_=1, for *λ*_*sim*_=2 the algorithm yielded *λ*_*est*_=4, and for *λ*_*sim*_=4 the algorithm yielded *λ*_*est*_=1. Importantly, for *λ*_*sim*_=1, the algorithm scored the ground truth (*λ*=2) in the same order of magnitude as the best estimation [Fig. 13]. This slight discrepancy underscores the importance of considering other *λ* values with a score comparable to the best estimation, as they may represent an acceptable solution. Thus, we encourage users to treat the algorithm output as an estimation for the range of probable λ values.

**FIG. 5.**
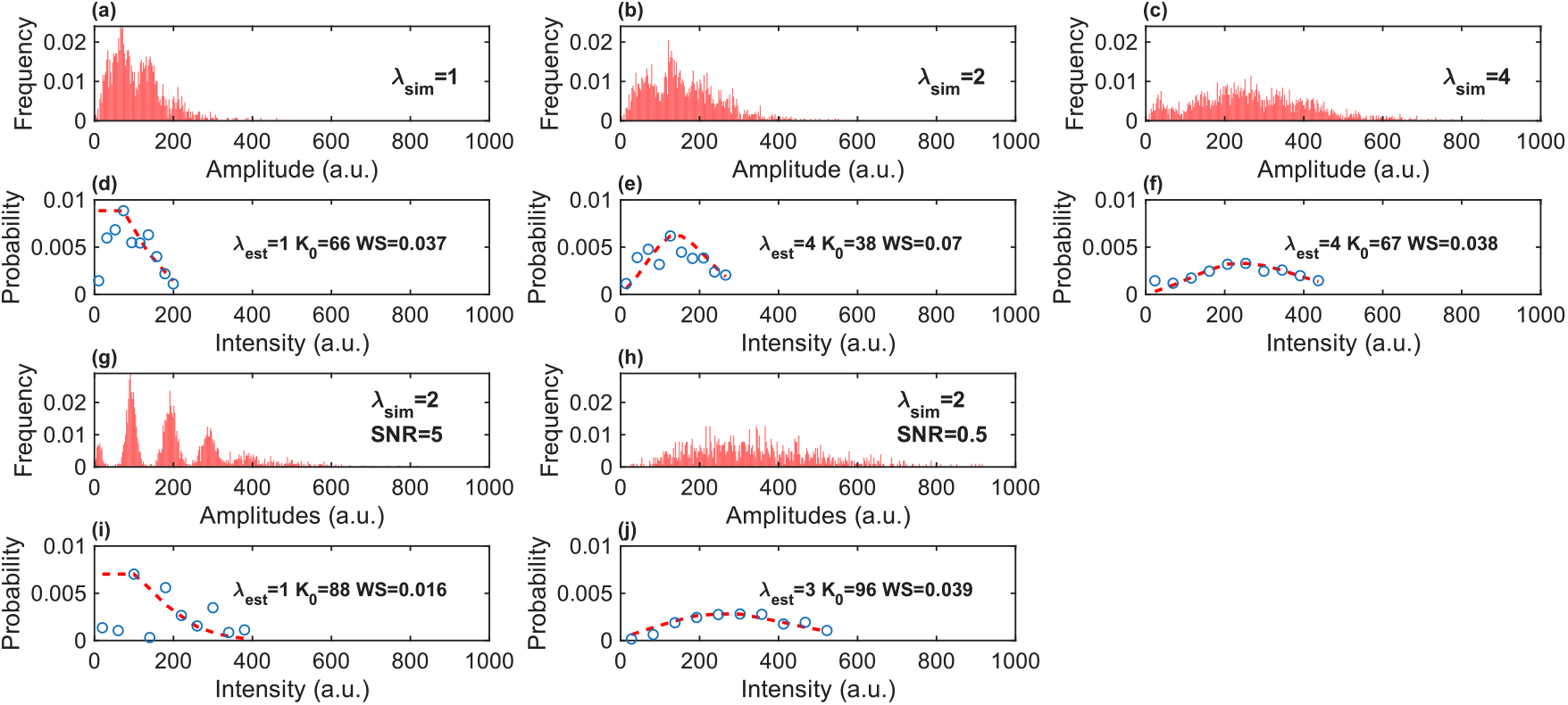
Estimating simulation parameters. (a-c) Amplitude distributions gathered from 1000 bursty simulated signals with similar base intensity (*k*_*0*_) and different Poisson parameter (*λ*) values. (a) *λ*_*sim*_=1 (b) *λ*_*sim*_=2 (c) *λ*_*sim*_=4. The distributions demonstrate a visible shift in accordance with the Poisson parameter used in the simulation. (d-f) Poisson distribution fits of the value used in the simulation. Blue points show the measured amplitude distribution, and red lines are the theoretical Poisson distribution. The fits show a relatively accurate estimation of the base intensity value *k*_*0*_. (g-h) Amplitude distributions gathered from 1000 bursty simulated signals with different SNR conditions. (g) SNR=0.5, (h) SNR=5. (i-j) Poisson distribution fits of the best estimates according to the algorithm in terms of *WS*. In the high SNR case, the estimation fails due to over-truncation. In the low SNR case, truncation is useful in providing correct estimates.

We further tested the estimation performance under different signal to noise ratios (SNRs) by simulating two more sets of signals. The first set had *λ*_*sim*_=2, *k*_*0*_=200, and a white Gaussian noise with a peak-to-peak amplitude of 40 (a.u.), corresponding to an SNR of 5. The second set had *λ*_*sim*_=1, *k*_*0*_=60, and white Gaussian noise with a peak-to-peak amplitude of 120 (a.u.), corresponding to an SNR of 0.5. We estimated the Poisson parameters of the amplitude data gathered from these signals [Figs. 5(g)-5(j)]. For both cases, the algorithm yielded *λ*_*est*_ values within the range [*λ*_*sim*_+1, *λ*_*sim*_-1]. In the high SNR case, our algorithm estimated *λ*_*est*_=1 and *k*_*0,est*_=88. We noticed that the algorithm correctly estimates the true *λ*_*sim*_ when we do not truncate the data. This is likely since in high SNR data, small amplitude signals are becoming harder to neglect. This potential issue should not affect real biological data that is intrinsically noisier. For the low SNR case, the algorithm estimated *λ*_*est*_=3 and *k*_*0est*_=96. These findings illustrate that one must be careful when neglecting some percentile of the data: while truncation of noisy, biological signals is acceptable, samples with a high SNR may require no truncation.

## V. EXPERIMENTAL VALIDATION

We validated our algorithm on experimental data from RNA-protein (RNP) granules formed *in vivo* within bacterial cells, and *in vitro* using ambient lab conditions. The granules were produced using our previously established phase separation system, which consists of synthetic RNA molecules (slncRNA) that fold into secondary structures with multiple hairpins, together with phage coat proteins capable of binding these hairpins. RNA molecules, bound by one or more proteins, intermittently enter or exit the granules, causing fluctuations in fluorescence levels, with the amplitudes of these shifts being dependent on the number of proteins bound to the RNA [18].

### A. GelMetrics reveals a simple Poisson distribution of single RNA bursts

We prepared three sets of granules *in vitro*, with a constant RNA concentration (120 nM) and varying protein concentrations, resulting in 1:1, 10:1 and 100:1 protein-to-RNA ratios (see Methods). The RNA construct, designated as PCP-14x/MCP-15x, contains 14 hairpin binding sites for the PP7 phage coat protein (PCP), interspaced by 15 hairpin binding sites for the MS2 phage coat protein (MCP), which serve as structural spacers and are not recognized by PCP [19]. This experimental setup enabled us to establish a scale for evaluating the algorithm’s performance, as we anticipated a similar Poisson parameter *λ*, but different *k*_*0*_ values (see Eq. 1.9), depending on the protein content in the system. We tracked the fluorescence intensity of multiple granules over a span of 60 minutes, with measurements taken every ten seconds, and applied our algorithm. The results show that regardless of protein content, most burst amplitudes appear near the lower end of the spectrum [0-100 (a.u.)], suggesting that free-floating proteins or RNA molecules bound by a few proteins traverse the phase boundary more easily [Figs. 6(a)-(c)]. We estimated the Poisson parameters of the negative amplitude distributions, as these represent molecules which exit the granule and therefore are affected by its internal dynamics. In addition, we filtered the amplitudes to remove the top 10% absolute values of the data, as these are statistically sparse rare events or outliers which originate from technical difficulties in the granule tracking process. In terms of *WS*, the algorithm estimated a *λ*_*est*_ value of 1 for all three experimental cases. The *k*_*0*_ value increased from 39 (a.u.) for granules formed with equal concentrations of RNA and protein, to 63 (a.u.) for granules formed with 10:1 protein to RNA ratio, to 139 (a.u.) for granules formed with 100:1 protein to RNA ratio. Interestingly, the increase in fluorescence seems to follow a logarithmic dependence from 1:1 to 10:1 vs. 100:1, consistent with Flory-Huggins theory expectations for liquid-liquid or liquid-gel phase separation, where the dense phase concentration depends weakly on total concentration. This is typically reflected in having more granules, each with a slightly denser composition, to account for entropic costs [Figs. 6(d)-(f)].

**FIG. 6.**
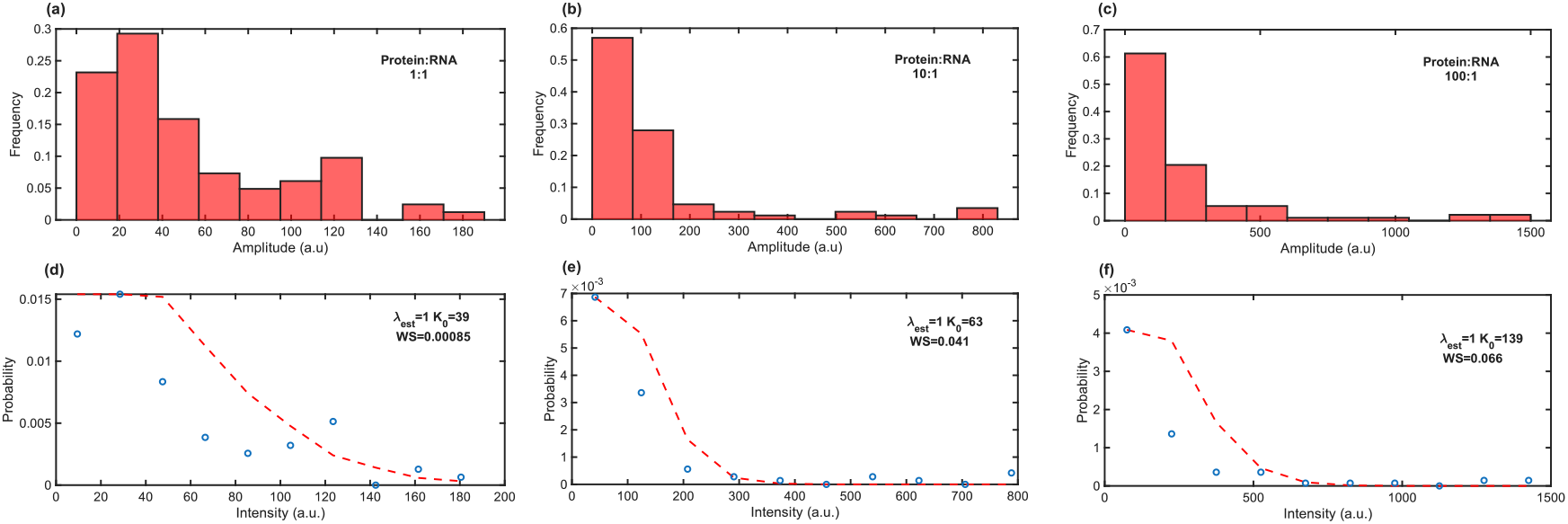
Analysis of *in vitro* experimental results. (a-c) Amplitude distributions gathered from RNP granules formed *in vitro*. In all three cases, reaction mixtures were prepared with an equal RNA concentration, but different protein concentration: (a) 1:1 protein to RNA ratio, ?, (c) 100:1 protein to RNA ratio, yielding a scale by which we can measure the performance of the algorithm. The amplitude distributions demonstrate a bias to the lower end of the scale, indicating that RNA molecules bound by few proteins can more easily traverse the phase boundary. (d-f) Poisson distribution fits of the best estimate according to the weighted score (*WS*). Blue points show the measured amplitudes distribution, red lines are the theoretical Poisson distribution. The fits show similar Poisson behavior (*λ*_*est*_=*1*), and a logarithmic increase in base intensity (*k*_*0*_). Plots (a), (b), and (c) represent measurements of 91, 96, and 103 negative amplitude bursts, respectively.

### B. QCP and PCP granules exhibit different material properties

We hypothesized based on our previous results [17,20] that a signal tracking algorithm can allow us to not only differentiate between granules with proteins containing an IDR element to ones without, but also extract internal material properties that can shed additional light on the underlying physical principles that guide this phase-separated system. To do so, we first tested our algorithm on experimental data from granules formed in *E. coli* cells. We utilized the Qβ and PP7 phage coat protein (QCP and PCP, respectively) together with three synthetic RNA sequences, one which encodes for ten QCP hairpin binding sites (designated QCP-10x), a second which encodes for five QCP binding sites and four PCP binding sites (designated QCP-5x/PCP-4x), and a third which encodes for 30 PCP hairpin binding sites (designated PCP-30x). QCP is an IDR containing phage coat protein, while PCP does not have this feature. Two plasmids, one encoding for the RNA molecules, and the other for either QCP or PCP fused to mCherry, were co-transformed into *E. coli* cells, which were grown overnight with full induction of the components.

We tracked the fluorescence intensity of the granules over 60 minutes, with measurements taken every 10 seconds, and applied our algorithm. We first tested our algorithm on PCP granules and confirmed our previously reported observations of a fluorescent signal interrupted by bursts composed of single RNA molecules, which are consistent with gel-like material properties. Our data shows that Poisson distributions characterized by *λ*=1 represent the most accurate description of our data for both QCP-5x/PCP-4x and PCP-30x [Figs. 7(a)-7(d)]. By contrast, analysis of the amplitude distribution of QCP [Fig. 8] shows a very different behavior, consistent with different material properties. The amplitude distributions of QCP granules for both RNA molecules present a profile more consistent with a Poisson distribution with *λ*>1 and appear to be rather similar in range, suggesting that QCP-10x RNA molecules which are not fully occupied more easily traverse the phase boundary [Figs. 8(a) and 8(b)]. Analysis of the Poisson estimations [Figs. 8(c) and 8(d)] reveal that in terms of *WS*, the distributions fit a Poisson with *λ*_*est*_=4, and *k*_*0*_=90 and 110, for QCP-5x/PCP-4x, and QCP-10x, respectively. This result is consistent with an interpretation of a brittle solid behavior, where the granules seem to periodically shed large chunks of proteins and RNA consisting of ~4 slncRNA molecules, reminiscent of spontaneous fracturing or fissioning of brittle solids or glasses (e.g., glaciers) [21].

**FIG. 7.**
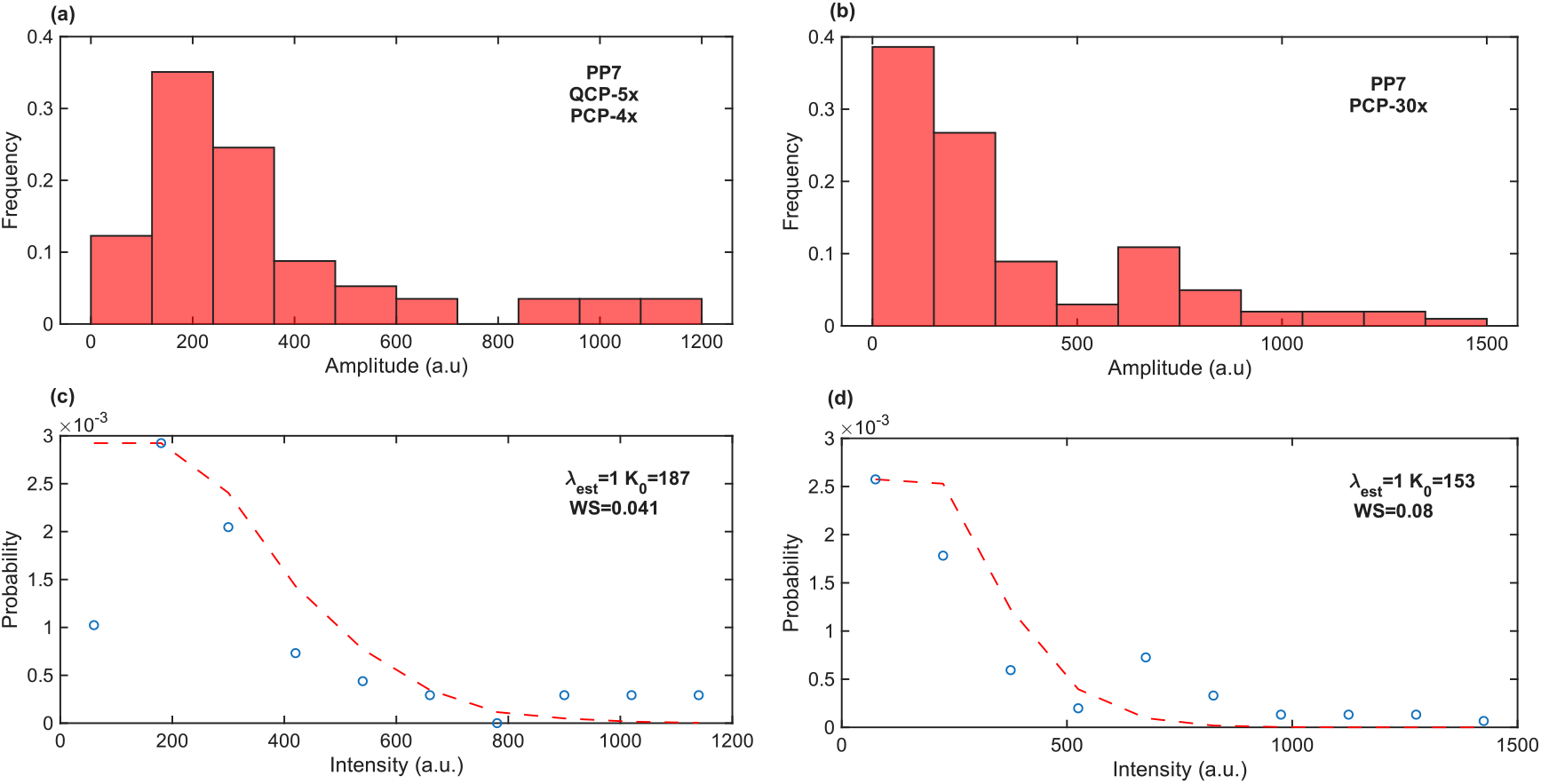
Analysis of PP7 *in vivo* experimental results. (a,b) Amplitude distributions gathered from RNP granules formed *in vivo* using PP7 and the RNA sequences (a) QCP-5x/PCP-4x, which has four binding sites for PP7, and (b) PCP-30x, which has 30 binding sites for PP7. (c,d) Poisson distribution fits of the best estimate according to the weighted score (*WS*). Here, the distributions best fit a Poisson function with *λ*=1. A larger *k*_0_ value is fitted to the higher valency RNA compared to the lower valency RNA case, as expected. However, the similarity in magnitude suggests PP7-30x is not fully occupied. Plots (a) and (b) represent measurements of 63 and 112 negative amplitude bursts, respectively.

**FIG. 8.**
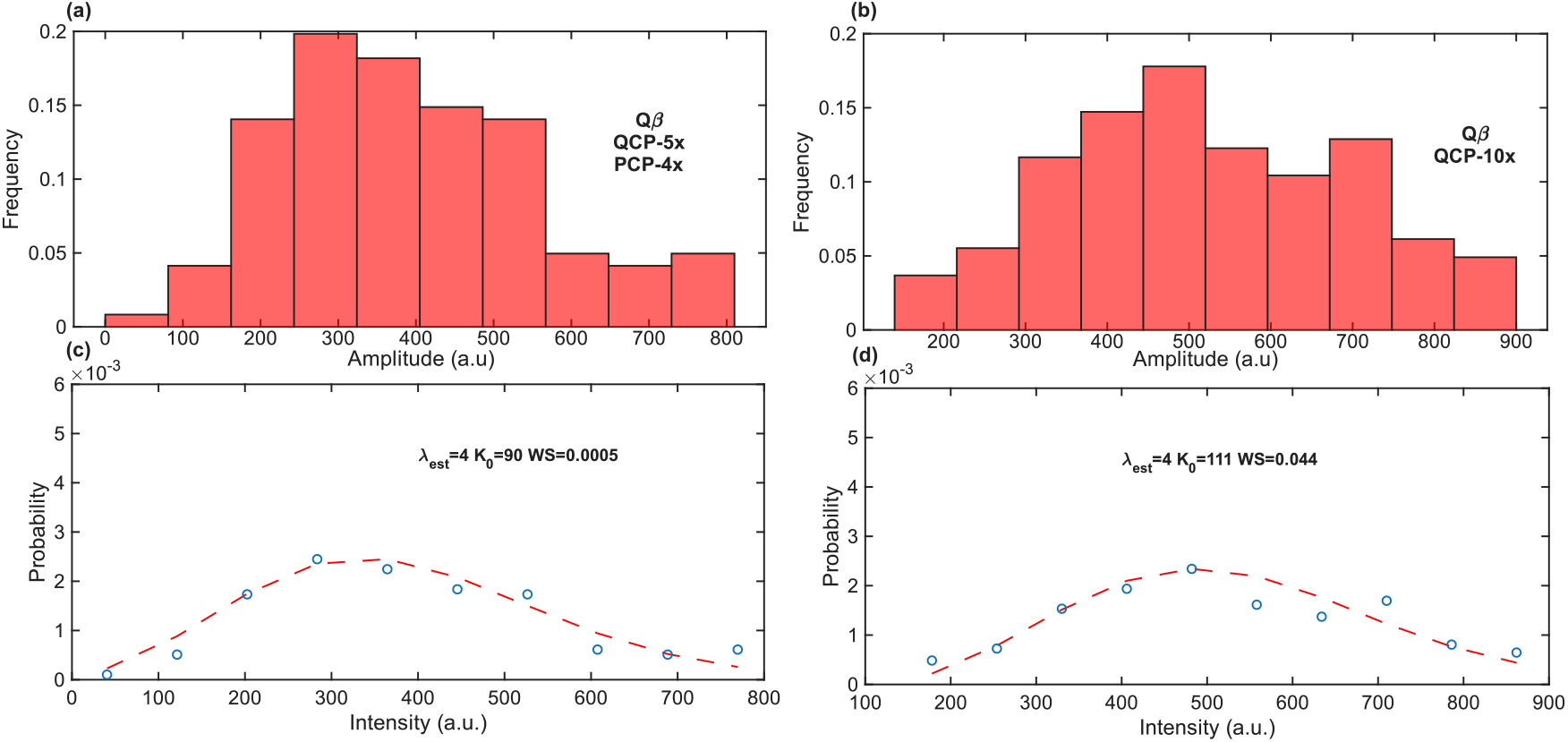
Analysis of Qβ *in vivo* experimental results. (a,b) Amplitude distributions gathered from RNP granules formed *in vivo* using Qβ and two different RNA sequences (a) QCP-5x/PCP-4x, which has five binding sites for Qβ, and (b) QCP-10x, which has ten binding sites for Qβ. (c,d) Poisson distribution fits of the best estimate according to the weighted score (*WS*). Here, the distributions best fit a Poisson function with *λ*=4 and with close *k*_*0*_ values, indicating that the QCP-10x RNA is likely not fully occupied by proteins. Plots (a) and (b) represent measurements of 134 and 181 negative amplitude bursts, respectively.

### C. Brittle behavior is associated with presence of IDR element

Given our previous observation [20] that QCP exhibited a multiphasic phase-separation behavior as a function of the RNA valency, we wondered whether the multiphasic behavior and the brittleness reported here were associated with the presence of IDR or were a general characteristic of QCP. To test this hypothesis, we constructed a PP7 protein fused to the highly-disordered fused in sarcoma (FUS) moiety [22]. We repeated our analysis with granules containing this hybrid IDR-construct (designated PP7-FUS). Our analysis shows that for the low-valency RNA cassette (QCP-5x/PCP-4x), the FUS addition shifts the most likely Poisson *λ*_*est*_ parameter to higher values that range from 1 to 3, and to lower *k*_*0*_ values, as compared with native PP7. This is consistent with our hypothesis that the disordered FUS domain leads to a potentially denser and brittle solid or glass-like granule phase [Figs. 9(b) and 16]. By contrast, for the high-valency cassette (PCP-30x), the burst signal analysis suggested no apparent effect for the FUS addition [Fig. 9(c) and 16], indicating that effect of the IDR moiety manifests itself only for low slncRNA valencies.

**FIG. 9.**
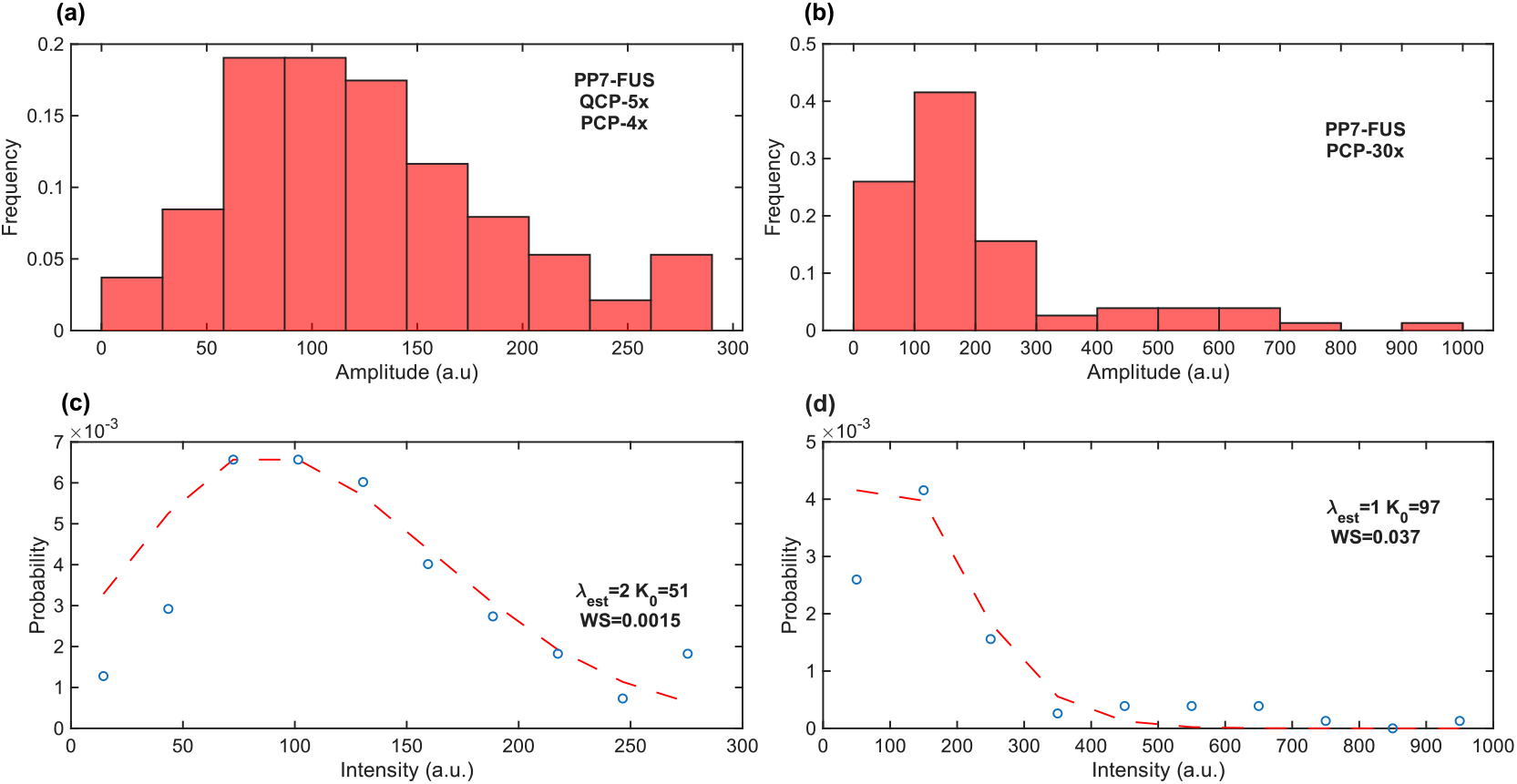
Analysis of PP7-FUS *in vivo* experimental results. (a,b) Amplitude distributions gathered from RNP granules formed *in vivo* using PP7-FUS and the RNA sequences (a) QCP-5x/PCP-4x, which has four binding sites for PP7-FUS, and (b) PCP-30x, which has 30 binding sites for PP7-FUS. (c,d) Poisson distribution fits of the best estimate according to the weighted score (*WS*). Here, the distributions best fit a Poisson function with *λ*=2 and *λ*=1 for the low- and high-valency RNA, respectively. This implies that granules formed with low-valency RNA phase-transition towards a more brittle phase due to IDR addition, while granules formed with high-valency RNA do not. The similar-in-magnitude *k*_*0*_ values indicate that the PCP-30x RNA is likely not fully occupied by proteins. Plots (a) and (b) represent measurements of 210 and 86 negative amplitude bursts, respectively.

## VI. DISCUSSION

In this work, we describe and assess a method for investigating the dynamics of solid-like phase separated condensates. Various experimental assays are available to researchers studying phase separation and membraneless organelles. However, these are often inadequate for the case of solid-like condensates, where dynamic motion exists at a much slower rate in comparison to liquid-like bodies. Our algorithm leverages straightforward fluorescence microscopy measurements of the condensates and employs statistical approaches to identify signal segments likely corresponding to molecules entering or exiting the condensate. By aggregating the amplitudes of these segments collected from multiple granules, we derive a distribution, which we fit to a modified Poisson function characterized by two parameters: *λ*_*est*_, representing the average number of molecules traversing the granules’ phase boundary, and *k*_*0*_, denoting the average fluorescence intensity of a single molecule.

We validated the performance of our algorithm using three sets of experimentally formed granules. The first set consisted of granules formed *in vitro* under controlled conditions. In these experiments, the concentration of the phase-separating agent (RNA) was kept constant, while the concentration of the fluorescent protein was varied. Analysis of the signals recorded from these granules revealed that their dynamics correspond to a Poisson distribution with *λ*=1, consistent with gel-like material properties, regardless of protein concentration, indicating an average of one RNA-protein molecular complex traversing the phase boundary during a burst event. By contrast, the fluorescence of a single emitter (*k*_*0*_) increased logarithmically as function of increasing protein concentration. This suggests that the proteins are stored within the granules at increasing densities consistent with expectations from Flory-Huggins theory, which predicts that the densification emerges from overcoming entropic penalties.

The second set of experiments involved granules formed *in vivo*, comparing QCP and PCP granules. We demonstrated that granules composed of different proteins can be differentiated based on their burst behavior, likely reflecting varying structural and material properties. Specifically, for QCP granule dynamics fit a Poisson distribution with *λ*=4 that is consistent with a brittle solid-or gel-like body that occasionally fractures to smaller bodies. By contrast, PCP granules exhibited a Poisson distribution with λ=1, which is consistent with a fully interconnected heterotypic gel-like granule, as was also observed *in vitro* for this system. Finally, our third set of constructed consisted of a PCP fused to FUS, a known IDR element, to test if the brittle behavior is recovered. Our data confirms this conjecture, and thus provides a new physical element to the growing body of effects found in biocondensates that are caused or triggered by the presence of IDR moieties. Moreover, the result obtained for tdPCP-FUS granules matches well our previously proposed phase diagram for the Qβ granules [Fig. 10(a)]. Specifically, for low slncRNA valencies and proteins devoid of IDR elements, the granules seem to adopt a weakly interconnected gel-like structure that exhibit a relatively high fluorescent amplitude per burst. However, for low valency granules composed of proteins with IDR elements, a solid-like or glass-like phase seems to be adopted, characterized by increased *λ*_*est*_ and substantially smaller *k*_*0*_ value as compared with the non-IDR case [Fig. 10(b)] suggesting perhaps a denser structure where the protein emitters are partially quenched. Finally, for high-valency RNA molecules, the internal structure seems to return to a gel-like phase, independent of the presence or absence of IDR elements. In addition, when the fitted amplitude of the bursts is normalized by the slncRNA valency, the reduction observed in this value suggests that either the slncRNAs are not fully occupied by proteins at high valency, or that the granules themselves are denser, leading to fluorescent quenching.

**FIG. 10.**
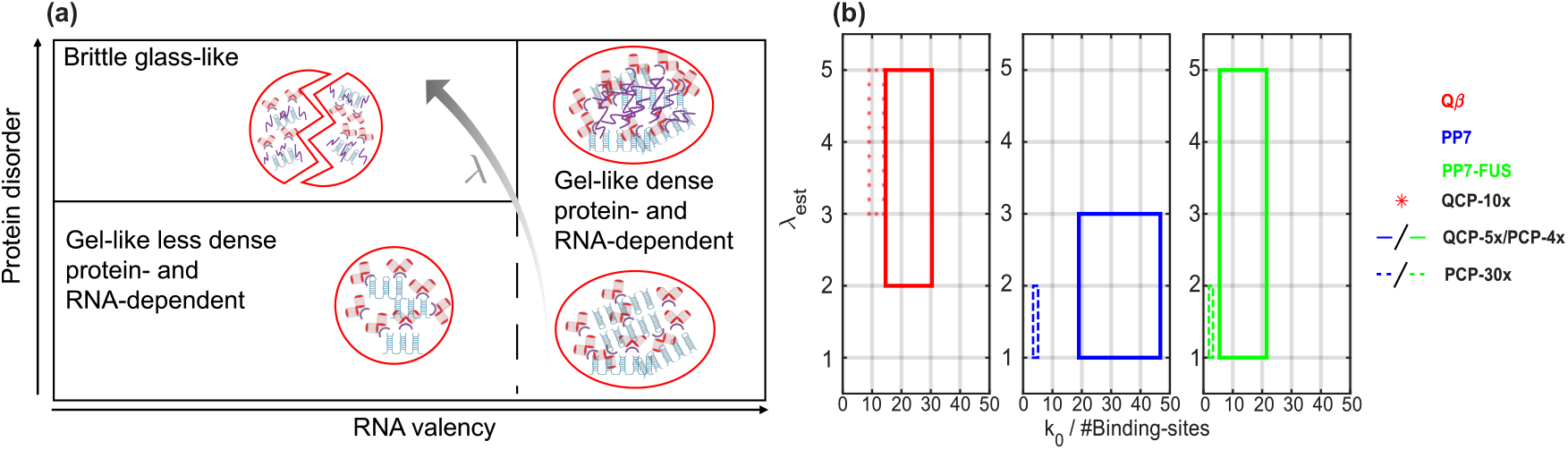
IDR shifts granule phase behavior towards brittle regime. (a) Proposed phase diagram for PP7 RNP granules. With low-valency RNA, FUS-absent PP7 molecules form gel-like granules, characterized by Poisson parameter *λ*=1. However, when the FUS-fused PP7 protein is introduced with the same RNA, the granules display a more brittle behavior, with increased probable *λ* values. In contrast, with high-valency RNA, both PP7 and PP7-FUS form denser gel-like granules, characterized by Poisson parameter *λ*=1 and large *k*_*0*_ values. (b) *λ* and *k*_*0*_ values obtained for granules formed with Qβ (red), PP7 (blue), or PP7-FUS (green) proteins, and with the QCP-5x/PCP-4x (solid line), QCP-10x (dotted line), or PCP-30x (dashed line) RNA. The addition of an IDR to the PP7 protein increases *λ* values for the low valency RNA case only.

Taken together, the results of our analysis suggest that the differences in granule kinetics may stem from structural and material characteristics that differentiate granules with and without an IDR element. These findings could be relevant to several diseases (e.g., repeat-expansion disorders [23–26]) where a substantial increase in long non-coding RNA (lncRNA) valency has been associated with pathology. Specifically, a transition between a brittle phase, that periodically releases large chunks of RNA and proteins into the cellular milieu, to a dense and strongly interconnected gel-like phase, that does not, can lead to a sharp reduction in the concentration of granule-bound protein(s) in the dilute cellular phase outside of the granules. Such a “step-like” knock-down of the supply of dilute-phase protein concentration may act similarly to a deletion mutation, which in turn may lead to a strong phenotypic effect.

The presented algorithm has ample room for further development. For example, incorporating machine learning methods to better identify signal segments corresponding to biological events, and adding regularization to the Poisson estimation process to place selective emphasis on the tail end of the distribution, which we noticed might bias the results. We expect this method will be useful in cases where different sets of granules need to be compared to one another. Such a scenario is typically encountered in perturbation experiments (mutations/knockouts of components) and in *in vitro* reconstitution studies, where components can be selectively added or removed to probe behaviour. Finally, while initially designed for analysis of the kinetics of solid-like condensates, we anticipate that with sufficient temporal resolution and adjustment of internal parameters, the algorithm could also be applied to liquid-like condensates, adding to the array of methods for condensate studies that exist today.

## VII. CONCLUSIONS

In conclusion, we present here a method of investigating non-liquid like phase separated condensates using standard fluorescence microscopy and achieving single molecule resolution. The reported parameters (*λ, k*_*0*_) support a more nuanced picture of the dynamic and structural nature of such condensates, and reveal a more complex phase diagram for RNA-protein granules, consisting of three distinct phases: a weakly interconnected, less-dense gel-like phase, a brittle solid-like or gel-like phase, and a likely strongly-interconnected and protein-dense gel-like phase.

## VIII. METHODS

### A. Formation of *in vitro* RNA-protein granules

Construction of the required plasmids for RNA and protein expression, and expression/synthesis of the components was done as previously described in [18]. Briefly, a vector containing the DNA sequence encoding for the required RNA was linearized via digestion with a restriction enzyme, followed by an *in vitro* transcription reaction with a HiScribe T7 RNA synthesis kit (New England Biolabs (NEB) #E2040). Following *in vitro* transcription, the remaining DNA in the reaction was degraded using a DNAse I enzyme (NEB #M0303S). RNA products were purified using Monarch RNA Cleanup Kit (NEB, #T2040S) and either used immediately or stored at −80 °C for later use.

For protein purification, *E. coli* cells expressing the fusion protein were grown overnight in 5 ml Luria Broth (LB) with appropriate antibiotics at 37 °C with 250 rpm shaking, and diluted the following morning 1/100 into Terrific Broth (TB: 24 g yeast extract, 20 g tryptone, 4 ml glycerol in 1 L of water, supplemented with 17 mM KH_2_PO_4_ and 72 mM K_2_HPO_4_), with antibiotic and inducer. Cells were grown at 37 °C and 250 rpm until they reached an optical density larger than 10. Cells were harvested, resuspended in 30 ml resuspension buffer (50 mM Tris-HCl pH 7.0, 100 mM NaCl and 0.02% NaN_3_), and disrupted by four passages through an EmulsiFlex-C3 homogenizer (Avestin Inc., Ottawa, Canada). Cell lysate was centrifuged at 10,000 g for 30 min to separate the protein content from cell debris. The fusion protein was purified using HisLink Protein purification resin (Promega, Madison, WI #V8821) according to the manufacturer’s instructions. Buffer was exchanged to 1xPBS (Biological Industries, Israel) using Amicon ultra-2 centrifugal filters (Merck, Burlington, MA #UFC203024) using the manufacturer’s instructions. *In vitro* experiments were performed in granule buffer (750 mM NaCl, 1 mM MgCl_2_, 10% PEG4000).

Reactions were set up and allowed to rest at room temperature for 1 h. 3–5 µl from the reaction was then deposited on a glass slide prior to imaging.

### B. Formation of *in vivo* RNA-protein granules

BL21-DE3 cells expressing one plasmid encoding for the slncRNA, and a second plasmid containing the fluorescent protein fused to a phage coat protein, were grown overnight in 5 ml LB at 37 °C, with appropriate antibiotics (Cm, Amp), and in the presence of two inducers: 1 mM IPTG (Ornat, # 67630), and 60 μM C4-HSL (Cayman Chemicals, #10007898). Overnight culture was diluted 1:50 or 1:25 into 3 ml semi-poor medium consisting of 95% bioassay buffer (BA, for 1 L: 0.5 g Tryptone [Bacto], 0.3 ml glycerol, 5.8 g NaCl, 50 ml 1 M MgSO_4_, 1 ml 10×PBS buffer pH 7.4, 950 ml DDW) and 5% LB with appropriate antibiotics and induction. Culture was shaken for 3 more hours at 37 °C. Bacterial immobilization was done by depositing 1.5 μl of the cell culture on a gel slide (3 ml PBSx1, mixed with 0.045 g SeaPlaque low melting Agarose (Lonza, Switzerland), heated for 20 seconds and allowed to solidify between two glass slides), prior to imaging.

### C. Fluorescence microscopy

*In vitro* granules and live *E. coli* cells were imaged in a Nikon Eclipse Ti-E epifluorescent microscope (Nikon, Japan) with a 100×1.45 NA oil immersion objective and using an Andor iXon Ultra EMCCD camera at 6 frames-per-minute with a 250 msec exposure time per frame. Excitation was performed by a CooLED (Andover, UK) PE excitation system at 585 nm for the mCherry protein.

### D. Image analysis and signal extrapolation

The top 10% of bright spots in the field of view were tracked using the ImageJ MosaicSuite plugin [27–29]. A field of view typically contained dozens of granules (*in vitro*) or cells containing puncta (*in vivo*). The tracking data (x,y,t coordinates of the bright spot centroids), together with the raw microscopy images, were processed by a custom Matlab (The Mathworks, Natick, MA) script, designed to extract granule and background signals from the raw data. For each bright spot, a 14-pixel wide sub-frame was extracted from the field of view, with the bright spot at its center. Each pixel in the sub-frame was classified to one of three categories, as follows. The brightest pixels were classified as ‘spot pixels’, typically appearing as a cluster which corresponds to the spot itself. The dimmest pixels were classified as ‘dark background’, representing empty regions in the field of view. Finally, unclassified pixels were then classified as ‘cell background’. From each sub-frame, two values were extracted: the mean of the ‘spot region’ pixels and the mean of the ‘cell background’ pixels, corresponding to spot intensity and background intensity values. This process was repeated for each spot and for each time point, resulting in sequences of intensity vs. time for both the spot and cell background.

## ACKNOWLEDGMENTS

This work was supported by the European Union’s Horizon 2020 Research and Innovation Programme under grant agreement no. 851615. This study was partially supported by the Israel Innovation Authority. We thank Dr. Nitsan Dahan and Dr. Yael Lupu-Haber from the Technion Life Sciences and Engineering Infrastructure Center for their support in microscopy experiments.

## APPENDIX A ALGORITHM PARAMETERS

As depicted in Eq. (1.10),we took into account multiple metrics to derive the final score. We noticed that for low simulated *λ* values, the algorithm yielded *λ*_*est*_ larger than anticipated [Fig. 11]. Inspection of the fitting process revealed the reason to be the presence of low-amplitude signals, leading to an intrinsic bias toward large *λ* values as the algorithm maximizes the fit regardless of the signal significance and is therefore prone to outliers. Thus, we decided to mitigate this effect by left-truncating our data, neglecting signals below the 15^th^ percentile. When truncation was performed, low *λ* values were correctly estimated. We also noticed that when taking into account only the MSE metric ([*w*_*MSE*_=1,*w*_*QQ*_=0, *w*_*KS*_=0]), some bias still remains. Therefore, to have the most accurate estimation of simulated data, we finally chose to work with weights of [*w*_*MSE*_=0.5,*w*_*QQ*_=0.25, *w*_*KS*_=0.25]

**FIG. 11.**
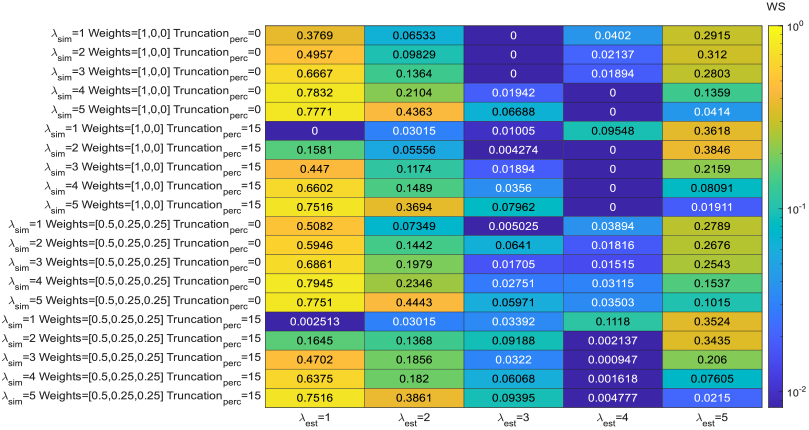
Algorithm parameters evaluation. The heatmap depicts weighted scores (*WS*) for every simulated and estimated Poisson parameter (*λ*_*sim*_ and *λ*_*est*_, respectively), as a function of different truncation percentiles (*Truncation*_*perc*_, data below this percentile was neglected), and different weights (*Weights*, in the order of [*w*_*MSE*_, *w*_*QQ*_, *w*_*KS*_]).

**FIG. 12.**
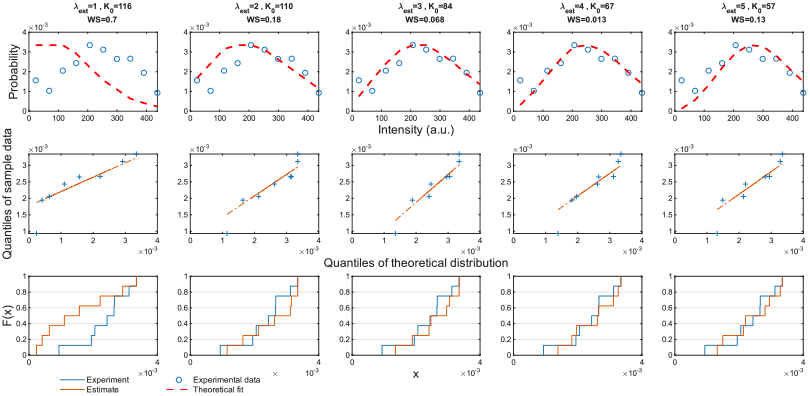
Sample fitting of an amplitude distribution to five modified Poisson functions, each with a different λ and *k*_*0*_ parameters. The top row depicts the experimental data (circles) and the theoretical fits (red lines). The center row depicts the QQ-plots of the fits. The bottom row depicts experimental and theoretical cumulative distribution function plots.

**FIG. 13.**
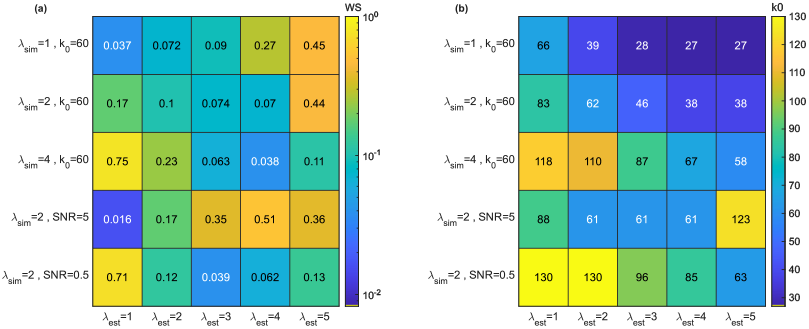
*WS* (a) and *k*_*0*_ (b) values obtained for simulated data, calculated for all *λ*_*est*_ values.

**FIG. 14.**
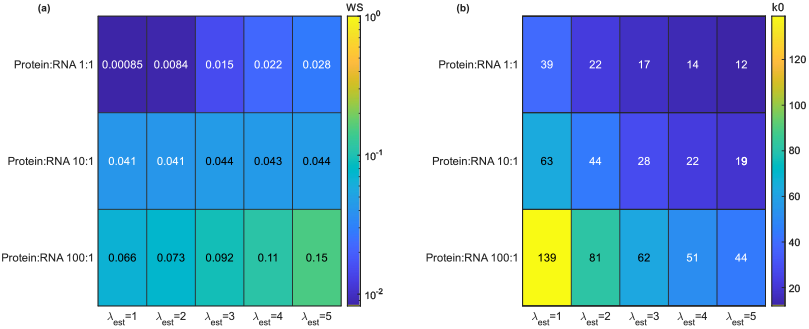
*WS* (a) and *k*_*0*_ (b) values obtained for *in vitro* experiments, calculated for all *λ*_*est*_ values.

**FIG. 15.**
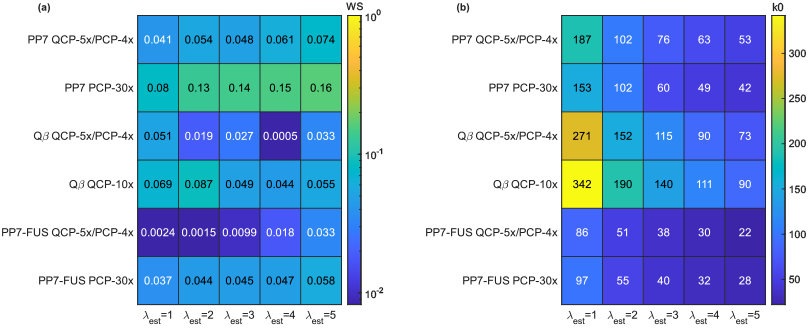
*WS* (a) and *k*_*0*_ (b) values obtained for *in vivo* experiments, calculated for all *λ*_*est*_ values.

**FIG. 16.**
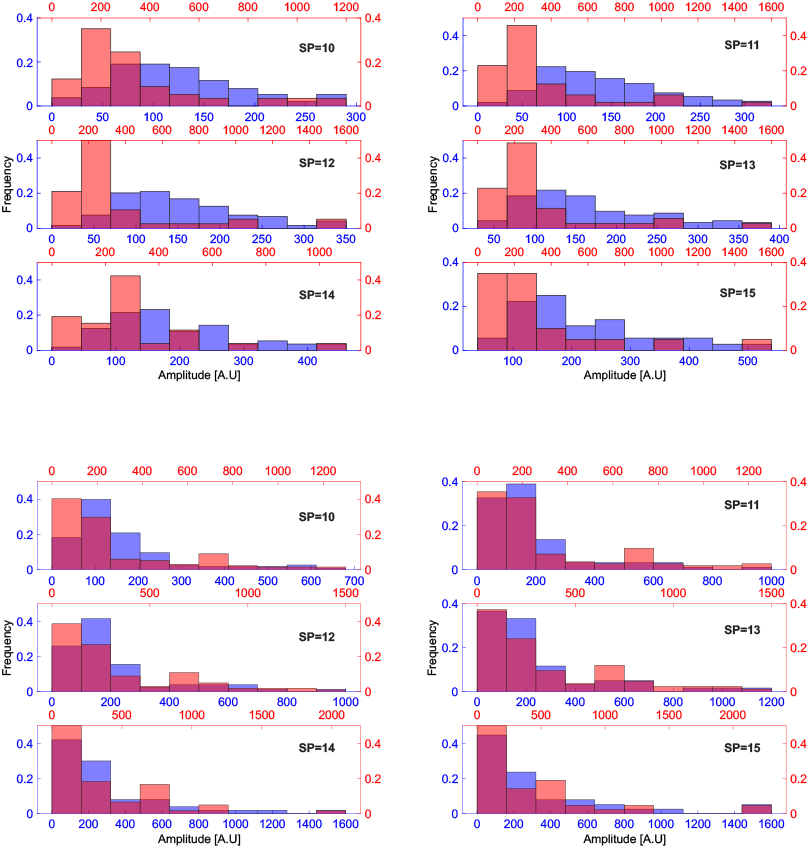
Experimental phase behaviors of granules made with PP7 (red) or PP7-FUS (blue) proteins and RNA encoding four (top) or 30 (bottom) PCP binding sites. The low-valency shows a histogram-peak shift while the high-valency RNA does not. Behaviors are consistent with a range of stringency parameters (SP).

## APPENDIX B TROUBLESHOOTING

We are aware that different experimental results, obtained by different instrumentations, might yield fits that seem to be poor. Thus, we suggest a few guidelines to help users to achieve the best outcomes. First and foremost, fits must pass the “eye-test”. During our testing of the algorithm, we noticed some fits got scored rather well despite an obvious deviation from the actual experimental data. We therefore recommend the user assess calculated fits qualitatively first, and if the fit appears of low-quality, adjust fit parameters. The first course of action to be taken is to change the stringency parameter (SP) used. Since SP dictates the number of events detected, a value too high may lead to too few experimental events for which to find a reliable fit. We recommend that the number of events detected be at least ~60. The SP can be increased for data for which many events were detected, thus lowering the noise from the real data. As a general rule, SP should not be lowered to values smaller than 10. Another feature that can be adjusted is the threshold percentile of the left-truncation of the data. As a default, the burst data is grouped into ten bins, and data below the 15^th^ percentile of said bins is neglected. We noticed that for data with unusually intense signals, such as the case for very high-valency RNA data, the bin width increases significantly. As a result, the neglected bins may contain too much “real” data. To address this issue, the user may use two separate approaches. The first is to change the number of bins the burst data is being grouped into. However, from our experience, this approach is usually extremely data-dependent and requires a distinct modification for each data set. A superior approach is to lower the threshold percentile for the truncation. We found the 10^th^ percentile to capture enough data to yield high-quality fits, while still leaving out the low intensity, negligible signals. This strategy was utilized to fit samples with high-intensity signals [Figs. 6(b), 6(c), 7(b), and 9(b)]. We strongly advise users to be cautious when lowering thresholds or SP, as these actions introduce further noise. One should consider compensating this effect by lowering one parameter (e.g., truncation threshold) while increasing the other (e.g., SP). The last parameter that can be tweaked is the weight given to each fit metric used by our algorithm. We generally suggest that users not alter these weights, as we looked for the most global settings, adequate for different sets of data.

## APPENDIX C FULL ANALYSIS RESULTS

To allow a thorough assessment of the algorithm, we present a full example of its output and report complete results obtained from simulated, *in vitro*, and *in vivo* data.

## DATA AVAILABILITY

The GelMetrics algorithm source code, together with *in vitro* and *in vivo* experimental data, are available at [30].

